# Genome-informed integrative taxonomic description of three cryptic species in the earthworm genus Carpetania (Oligochaeta, Hormogastridae)

**DOI:** 10.1101/802017

**Authors:** Daniel Fernández Marchán, Rosa Fernández, Jorge Domínguez, Darío J. Díaz Cosín, Marta Novo

## Abstract

Research on cryptic species complexes has reached a consensus on the necessity of integrating multiple sources of evidence. Low-coverage genomic scan techniques like Genotyping-by-Sequencing (GBS) have proven useful to study these groups. Both integrative taxonomy and genome-wide single nucleotide polymorphism (SNP) data remain to be widely applied to earthworms, an animal group with widespread presence of cryptic diversity. The genus *Carpetania* (formerly the *Hormogaster elisae* species complex) was found to contain six deeply divergent genetic lineages and some inconspicuous morphological differentiation based in a handful of Sanger-sequenced markers. Marchán et al. (submitted) delimited three well supported species-level clades on the basis of a genomewide SNP dataset and geometric morphometric analyses, highlighting the necessity of a formal taxonomic description of these taxa. In this work, further analyses are applied to the SNP data and a thorough morphological study is performed in order to provide an integrative description of two new species and to redescribe *Carpetania elisae*. Species-specific SNPs are identified and used as diagnostic characters, and genome-wide and cytochrome oxidase C subunit 1 (COI) genetic distances are compared finding a strong correlation between them. The taxonomic description of these three cryptic species provides a useful tool to include them effectively in ecological studies and biodiversity conservation actions.

## Introduction

Since the advent of molecular phylogenetic techniques, discovery of cryptic species complexes showed a surge in scientific interest (Bickford et al., 2007; Pfenninger & Schwenk, 2007; Trontelj & Fišer, 2009; León, de León, & Nadler, 2010; Nygren, 2014), followed by a stage of novelties in species delimitation methodologies including different algorithms, genetic markers and integrative approaches (Pons et al., 2006; Z. Yang & Rannala, 2010; Puillandre, Lambert, Brouillet, & Achaz, 2012; Zhang, Kapli, Pavlidis, & Stamatakis, 2013; Ziheng Yang, 2015). Two main lessons can be gathered from those studies: to obtain robust species delimitation hypotheses, several information sources (molecular, morphological, ecological, ethological…) must be integrated (Queiroz & De Queiroz, 2007); and nuclear molecular markers are necessary to rule out the possibility of confounding deep mitochondrial lineages with proper cryptic species (Dupont, Porco, Symondson, & Roy, 2016).

Some works have already shown the potential of genome-wide single nucleotide polymorphism datasets (either generated by RADseq or GBS-Genotyping By Sequencing-) to provide rich phylogeographic and species delimitation information for cryptic species complexes (Garg et al., 2016; Brunet et al., 2017, Rancilhac et al., 2019), but have only been applied to earthworms in the *Lumbricus rubellus* complex (Giska, Sechi, & Babik, 2015; Anderson, Cunha, Sechi, Kille, & Spurgeon, 2017). The first work failed to detect differentiation between sympatric cryptic lineages, which was interpreted as these lineages not corresponding to biological species. The second work, performed on different lineages of the same complex, found strong differentiation between them using similar analyses.

Pervasive cryptic diversity has been found in earthworms across different families (Lumbricidae - King, Andrew King, Tibble, & Symondson, 2008; Fernández, Almodóvar, Novo, Simancas, & Díaz Cosín, 2012; Shekhovtsov, Golovanova, & Peltek, 2013; Porco et al., 2018; Hormogastridae - Novo, Almodóvar, Fernández, Trigo, & Díaz Cosín, 2010; Megascolecidae - Chang, Lin, & Chen, 2008; Buckley et al., 2011, Moniligastridae - Ganin & Atopkin, 2018). Integrative taxonomy has yet to be widely employed in these complexes (Taheri et al., 2018).

One of the most studied cryptic species groups among these animals is the former *Hormogaster elisae* Álvarez, 1977, recognized as the genus *Carpetania* after Daniel Fernández Marchán et al., (2018). Six highly divergent cryptic lineages were identified using Sanger-sequenced mitochondrial and nuclear markers (Daniel F. Marchán, Fernández, de Sosa, Díaz Cosín, & Novo, 2017), but their description was precluded by the absence of clear-cut limits between the putative species. Quantitative differences in the distal end of genital chaetae were discovered between those cryptic lineages through geometric morphometrics (Daniel F. Marchán, Sánchez, et al., 2016), hinting a possible pseudocryptic status for the identified lineages. Pseudocryptic species are those classified as cryptic due to the “inadequacy of the morphological analysis” (Knowlton, 1993) and can usually be distinguished after careful morphological analysis together with molecular data (Lajus, Sukhikh, & Alekseev, 2015).

Recently, a rich genome-wide SNP dataset was obtained through GBS for seventeen populations and 85 individuals of *Carpetania* by Marchán et al. (submitted), with the main objective of studying selection signatures and local adaptation in the cryptic complex. The authors applied different approaches to genetic structure identification and species delimitation together with geometric morphometrics analysis (Fig. 1), finding congruent support for three species-level genetic clusters (hereafter cluster A, B and C), which comprised one or more of the six previously identified lineages as described in Daniel F. Marchán et al., (2017).

**Figure 1.**
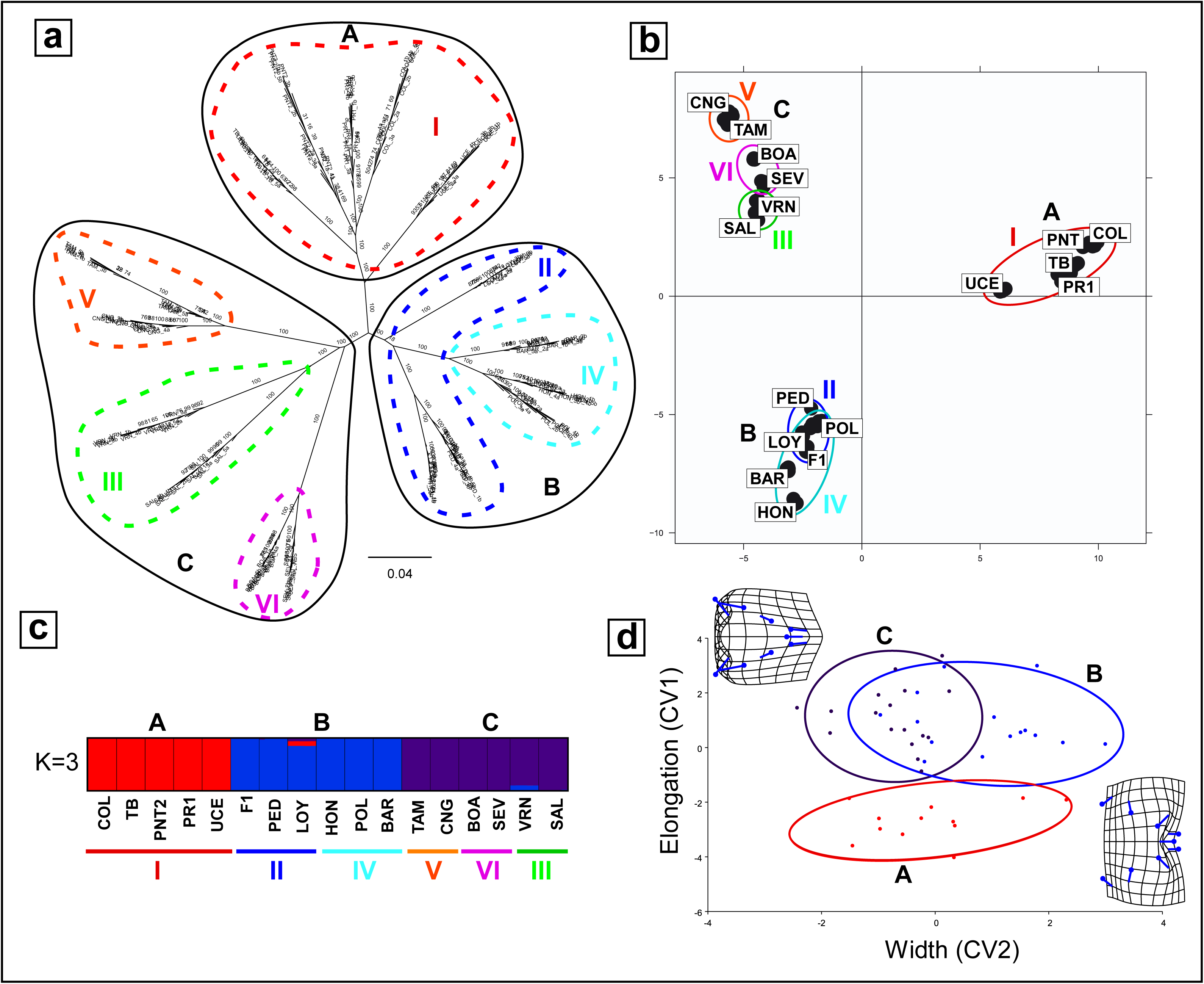
Different approaches to genetic structure identification and species delimitation in the *Carpetania* complex (modified from Marchán et al. submitted). Previously defined cryptic lineages (Daniel F. Marchán et al., 2017) are represented by a colour and roman number. Color codes and nomenclature are kept throughout the manuscript. a) Maximum likelihood inference of the phylogenetic relationships of the studied populations of *Carpetania*. Solid outlines indicate the species-level genetic clusters (A, B, C) found by the other analyses b) Principal Component Analysis (PCA) c) Barplot of STRUCTURE analysis. Each color shows percentage of assignment to a cluster or ancestral population. d) Canonical variate analysis of the shape of the genital chaetae of *Carpetania* main genetic clusters.

The well-supported putative species identified in *Carpetania* demanded a full, detailed taxonomic description: several authors have stressed the necessity of going a step further from cryptic species identification (Jörger & Schrödl, 2013; Wang et al., 2016). To this end, this work will build upon the framework provided by Marchán et al. (submitted), with the following objectives: i) identify genome-wide nucleotidic positions to be used as diagnostic characters in species description (molecular taxonomy); ii) explore further morphometric and internal anatomic characters to reinforce species description; iii) formally describe the species within *Carpetania*; iv) compare genomic and barcode genetic distances to validate the widespread use of the latter.

## Materials and methods

### Molecular data

SNP data comprising the geographical distribution and internal lineages of *Carpetania* was generated in Marchán et al. (submitted). In brief, GBS libraries were generated and sequenced for seventeen populations with five individuals from each totalling 85 individuals. Different datasets were generated with STACKS2 (Rochette, Rivera-Colón, & Catchen, n.d.) based on: *de novo* assembly/mapping to a reference transcriptome of C. elisae (‘reference-all SNPs’ dataset) and inclusion of all SNPs/one random SNP per locus (‘de novo-one SNP dataset).

‘de novo-one SNP’ and ‘reference-all SNPs’ datasets were further analysed in this work to estimate genetic diversity and to identify diagnostic positions, respectively.

Genetic diversity of the studied populations was described through identity by state (IBS) genetic distance within and between populations and fixation index (F_ST_). These parameters were obtained from the *populations* function (STACKS2 package) summary files. Correlation between SNP-based genetic distances and F_ST_ and the same parameters obtained from cytochrome C oxidase 1 sequences (Daniel F. Marchán et al., 2017) was tested through a Mantel test in the R package vegan (Dixon, 2003).

Diagnostic SNP positions were identified using function *nucDiag* in the R package spider (Brown et al., 2012) and parsed to the corresponding contig using the *Carpetania elisae* transcriptome after it was functionally annotated with eggNOG-mapper(Huerta-Cepas et al., 2017). Diagnostic nucleotidic positions were also identified in the molecular markers COI, 16S-tRNAs, 28S and H3 retrieved from (Daniel F. Marchán et al., 2017) following the same method.

### Morphological data

#### -External anatomy and morphometric characters

External anatomy characters commonly used in earthworm taxonomy were studied, with special attention to average weight and number of segments (found to have strong phylogenetic signal in Hormogastridae -Daniel F. Marchán, Novo, et al., 2016). These were measured in at least five mature individuals per population in a total of 22 populations, 9 populations from Cluster A, 6 populations from Cluster B and 7 populations from Cluster C. Statistical significance of differences was evaluated using ANOVA, Fisher’s LSD and Kruskal Wallis tests in Statgraphics 18.

#### -Internal anatomy

Internal anatomy characters commonly used in earthworm taxonomy were studied, with special attention to the only variable internal character in the *Carpetania* species complex, relative position of septum 9/10 and spermathecae. In this genus, septum 9/10 can appear displaced backwards (to 10/11) in its dorsal insertion, resulting in the spermathecae of segments 9 and 10 belonging functionally to the same segment, or show an unmodified disposition separating both pairs of spermathecae. This character was studied in 22 populations as well.

## Results and discussion

### -Genomic divergence and diagnostic positions

Genetic divergence parameters (average IBS distances and F_ST_ values) are represented in Fig. 2 (values can be found in suppl. Table 1). IBS distances between populations within the main clusters ranged from 0.09 to 0.15 (mean= 0.12) in both clusters A and B, while they ranged from 0.07 to 0.22 (mean=0.17) in cluster C. IBS distances between populations of different clusters ranged between 0.15-0.24 (mean=0.20) for clusters A and B, while they ranged between 0.18-0.24 (mean=0.21) for cluster C *vs* A and B.

**Table 1.** Diagnostic nucleotidic positions for the three species of the genus *Carpetania*. First nucleotide is the character state found on the species and the second nucleotide is the state for the rest of the species. Putative protein coded by the contig in where diagnostic positions are located is indicated when annotation is available.

| SNP | Scaffold |  | Base | Protein |
| --- | --- | --- | --- | --- |
| Cluster A |  |  |  |  |
| 344 | COMP190067_C0_SEQ1 | 395 | T/C |  |
| 727 | COMP206745_C0_SEQ1 | 527 | G/A | Retinol binding protein |
| 784 | COMP207582_C0_SEQ2 | 792 | G/C | SWAP switching B-cell complex 70kDa subunit |
| 949 | COMP209848_C1_SEQ1 | 269 | T/G |  |
| 1009 | COMP210899_C0_SEQ5 | 908 | A/T | Zinc finger, FYVE domain containing 20 |
| 1113 | COMP211847_C2_SEQ1 | 3393 | A/G | ATP-binding cassette, subfamily B (MDR TAP), member |
| 1291 | COMP213230_C0_SEQ4 | 658 | T/C |  |
| 1409 | COMP213929_C0_SEQ2 | 1505 | A/G | nuclear respiratory factor 1 |
| 1425 | COMP214029_C0_SEQ1 | 1951 | T/C | NADP-dependent oxidoreductase domain containing 1 |
| 1624 | COMP215394_C0_SEQ1 | 978 | T/C |  |
| 1784 | COMP216453_C0_SEQ4 | 1281 | G/C |  |
| 1814 | COMP216630_C0_SEQ7 | 1725 | C/T | leucine rich repeat and Ig domain containing 4 |
| 1842 | COMP216828_C1_SEQ13 | 4668 | G/C | Protein phosphatase 1 regulatory subunit 37 |
| 2127 | COMP218319_C1_SEQ44 | 2385 | T/C | Armadillo repeat containing 4 |
| 2169 | COMP218535_C0_SEQ2 | 1141 | C/G | Enzyme required for electron transfer from NADP to cytochrome P450 |
| 2267 | COMP219108_C2_SEQ1 | 3021 | A/G | SH3-domain binding protein 5 (BTK-associated) |
| 2417 | COMP219800_C0_SEQ6 | 2892 | A/C | CREB regulated transcription coactivator 1 |
| 2609 | COMP220400_C0_SEQ6 | 2547 | A/C | Nucleoporin 54kDa |
| 2639 | COMP220522_C0_SEQ13 | 4055 | G/T | ubiquitin- specific peptidase |
| 2652 | COMP220555_C0_SEQ8 | 3693 | T/C | integrator complex subunit 1 |
| 2712 | COMP220746_C1_SEQ15 | 359 | T/C | crumbs homolog 2 (Drosophila) |
| 2800 | COMP220908_C0_SEQ3 | 6612 | A/T | small nuclear RNA activating complex, polypeptide 4, 190kDa |
| 3071 | COMP62959_C0_SEQ1 | 45 | A/G |  |
| Cluster B |  |  |  |  |
| 195 | COMP1730661_C0_SEQ1 | 182 | T/G |  |
| 1410 | COMP213937_C2_SEQ1 | 662 | T/C | Solute carrier family 25 member |
| 1512 | COMP214730_C0_SEQ1 | 1067 | T/G | Peptidase family M41 |
| 2800 | COMP220908_C0_SEQ3 | 6612 | T/A | small nuclear RNA activating complex, polypeptide 4, 190kDa |
| Cluster C |  |  |  |  |
| 491 | COMP200542_C1_SEQ2 | 499 | A/C | Galactose binding lectin domain |
| 782 | COMP207582_C0_SEQ2 | 710 | G/T | SWAP switching B-cell complex 70kDa subunit |
| 893 | COMP209092_C1_SEQ1 | 734 | A/T | phosphodiesterase PDE8B |
| 1277 | COMP213187_C0_SEQ1 | 867 | G/A | Organic Anion Transporter Polypeptide (OATP) family |
| 2169 | COMP218535_C0_SEQ2 | 1141 | G/C | Enzyme required for electron transfer from NADP to cytochrome P450 |
| 2397 | COMP219681_C0_SEQ1 | 375 | G/A | PRP40 pre-mRNA processing factor 40 homolog |
| 2430 | COMP219875_C0_SEQ3 | 3810 | C/T | collagen, type XII, alpha 1 |
| 2776 | COMP220864_C4_SEQ4 | 4882 | G/A | kiaa2026 |

**Figure 2.**
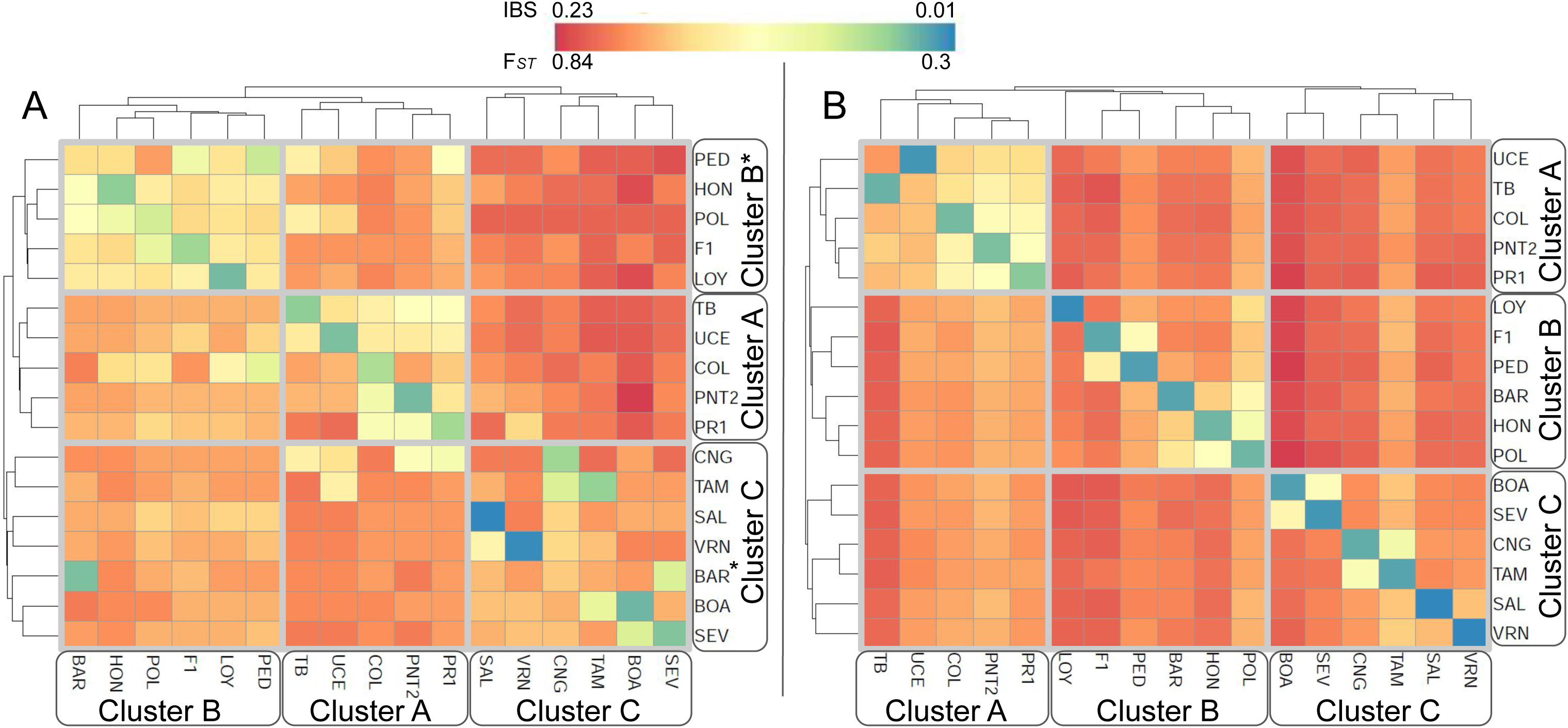
Heatmap and clustering graphs displaying FST values (left) and Identity by state (IBS) distances (right) between populations of *Carpetania* obtained from genome-wide SNPs.

F_ST_ values between populations within the main clusters were lower for cluster A (0.30-0.60, mean= 0.46) than for the other two clusters (B: 0.31-0.76, mean=0.59; C: 0.30-0.75, mean=0.63). F_ST_ values between populations of different clusters showed similar ranges for all of them (A: 0.51-0.82, mean= 0.65; B: 0.51-0.84, mean= 0.68; C: 0.55-0.79, mean= 0.67). Clustering analysis of both IBS distances and F_ST_ values (Fig. 2) recovered the same clusters as the phylogenetic inference and subsequent analyses.

Mantel test for genomic IBS distances and uncorrected pairwise COI distances showed a strong, statistically significant correlation between both sets of values (r = 0.7956, p = 0.001). Mantel test for genomic and COI-based F_ST_ values showed a weaker, statistically significant correlation between them (r =0.4801, p = 0.002).

These results suggest that divergence in COI sequence in the *Carpetania* species complex reflects to a significant extent genetic divergence across the whole genome, supporting the use of this molecular marker as a proxy for the identification of the cryptic species. COI barcoding (Hebert, Ratnasingham, & de Waard, 2003) has been widely accepted by the scientific community as a fast, simple and standardized method to identify, classify and delimit species (Decaëns, Porco, Rougerie, Brown, & James, 2013). It is worth noting that lineages identified on the basis of COI distance alone must be supported with additional molecular (nuclear markers), morphological, ecological, ethological and biogeographical evidence (Rougerie et al., 2009).

Several diagnostic SNP positions were identified: twenty-three for Cluster A, four for Cluster B and eight for Cluster C (table 1). Twenty-eight diagnostic SNP positions were assigned to the putative protein coded by the surrounding region of the variant nucleotide (table 1). Diagnostic SNP positions were found in genes with different biological functions, as regulation, transport, response to stimuli, developmental processes, cellular processes, multicellular organismal processes and metabolism among others.

A single diagnostic nucleotidic position was found for 28S molecular marker, distinguishing Cluster C from the other two. No diagnostic positions were identified for the rest of the Sanger-sequenced molecular markers.

Species-diagnostic SNPs have been successfully identified in plants (Cullingham, Cooke, Dang, & Coltman, 2013) and fishes (Hand et al., 2015). Even though they may appear more difficult to use for species identification than diagnostic positions in traditional Sanger-sequenced molecular markers, the cost of GBS and RAD-sequencing has lowered significantly making their use in taxonomic studies more viable. Another alternative is the development of SNP arrays based on the previously established (through GBS/RADseq) diagnostic SNPs, as an inexpensive, automatized approach.

### -Morphological analyses

Studied Cluster A and Cluster B populations showed significantly different average body weights (3.03 grams vs 4.98 grams) according to the different statistical tests. However, Cluster C average body weight (3.89 grams) was statistically indistinguishable from the other two.

No significant differences were found between the average number of segments of the studied populations of Clusters A, B and C, even though Cluster B showed the smallest average (248 segments) and C showed the highest (273 segments).

The difference in average values of morphometric characters is reminiscent of the ones found between *Lumbricus terrestris* and its sibling species *Lumbricus herculeus* (James et al., 2010). As found here, differences in weight and number of segments were not clear-cut, with overlapping distributions. This precludes the use of these characters in the diagnosis of the cryptic species, yet they are valuable for preliminary assignment in the field.

Character states for the relative position of spermathecae and septum 9/10 (separated/not separated by septum 9/10) showed high consistency for individuals from each population. However, those character states were not shared by all populations within the clusters (Fig. 3), with the exception of cluster B -where all studied populations showed spermathecae not separated by septum 9/10. Cluster A populations showed separated spermathecae, but the most basal populations within the clade showed not separated spermathecae. For cluster C this character showed no clear pattern, with two of its internal lineages showing constant states and the third containing populations with either character state. Both described species in the sister genus *Diazcosinia* possess spermathecae not separated by septum 9/10, suggesting this could be the ancestral character state for *Carpetania*.

**Figure 3.**
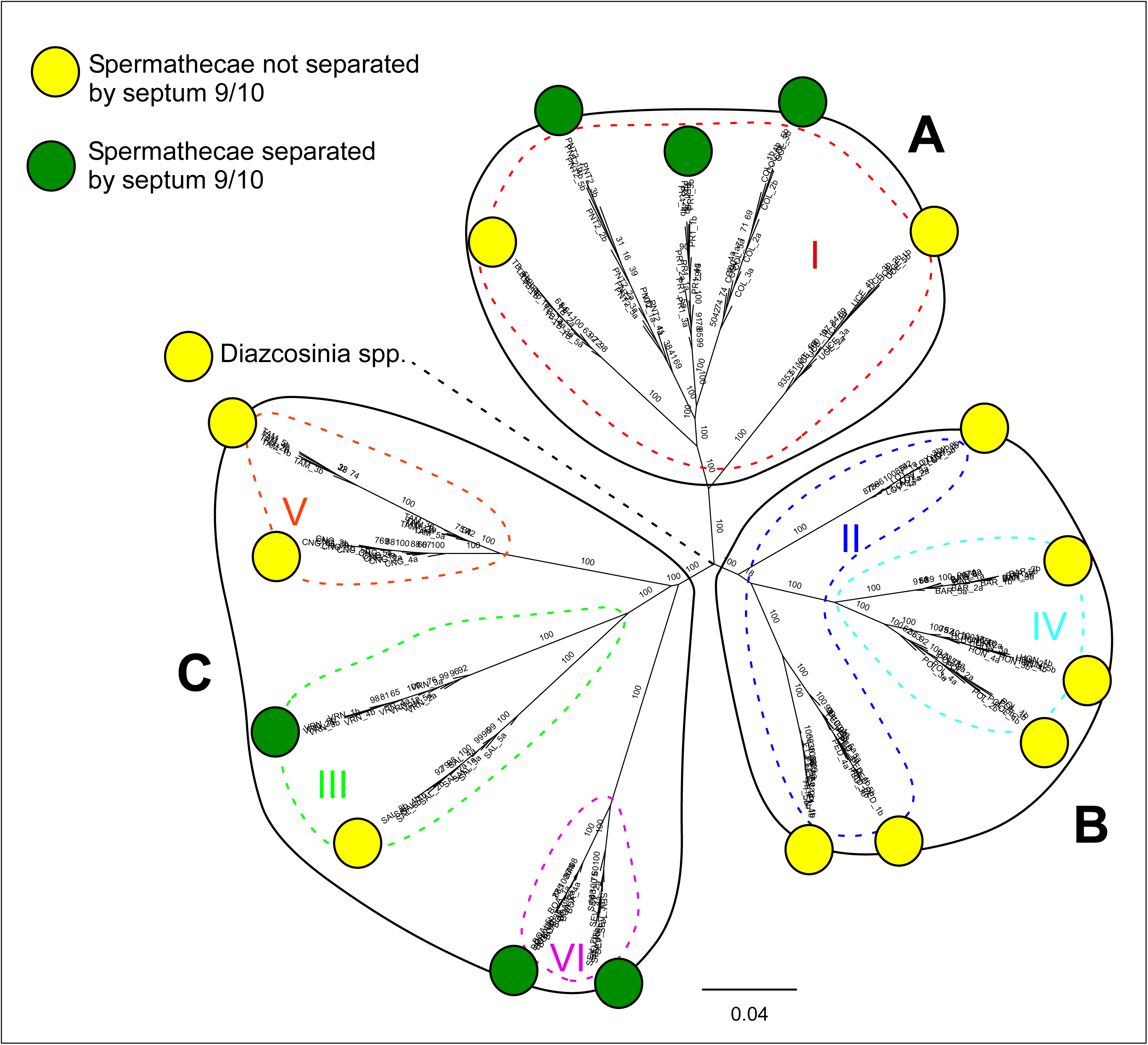
Relative position of spermathecae and septum 9/10 for the populations included in the phylogenetic analysis. A hypothetical position of the sister genus *Diazcosinia* (not included in the phylogenetic reconstruction) and its character state are shown by a dashed line. Lineages from (Daniel F. Marchán et al., 2017) are shown with the same code and colors (roman numbers, dashed outlines). Species-level clusters are represented by solid outlines and letters A, B, C.

The complex evolution of this trait disallows its use as diagnostic taxonomic character. However, it is an interesting example of multiple independent events of regression of a character state. Spermathecae not separated by septum 9/10 due to the backward displacement of the septum appears as a derived character state present in the common ancestor of *Carpetania* and *Diazcosinia*, while spermathecae separated by septum 9/10 is the most common disposition in other species of Hormogastridae and Lumbricidae. In order to disentangle the evolutionary pressures behind these changes, it would be necessary to understand the biological advantage of both character states. Even when the mechanisms of filling and release of sperm in earthworm spermathecae is not fully known, it is likely that contraction (and subsequent increased celomic pressure) of each segment should have a role in the process. Under this assumption, spermathecae separated by a septum would function independently, while spermathecae belonging functionally to the same segment would work coordinately. This would be relevant in the context of sexual selection and hermaphroditic sexual conflict: the capability of controlling which spermathecae stores sperm from a different mate is favourable to the female part, while diminished control over this uptake would be favourable to the male part. (Novo, Almodóvar, Fernández, Gutiérrez, & Díaz Cosín, 2010) found no evidence of differential sperm storage from different partners in each of the four spermathecae in *Carpetania* specimens from El Molar, which according to our hypothesis should have the ability to control front and back spermathecae separately. On the other hand, different sperm storage in front and back spermathecae has been observed in other earthworms (*Lumbricus terrestris* - Koene, Pförtner, & Michiels, 2005 -, *Eisenia andrei* - Porto, 2014). Further research would be needed to test this hypothesis and to propose other alternatives.

### -Taxonomic implications

The genetic structuring of *Carpetania* populations into three clearly separated clusters, characterized by significant genome-wide divergence, inconspicuous but detectable morphological differences and with no admixture between them calls for the formal taxonomic description of those clusters as species. Cluster B includes the type locality of *Hormogaster (Carpetania) elisae*, hence the original description is assigned to this clade. Meanwhile, Clusters A and C are described below as new species.

**Phylum Annelida Lamarck, 1802**

**Subphylum Clitellata Michaelsen, 1919**

**Class Oligochaeta Grube, 1850**

**Superorder Megadrili Benham, 1890**

**Order Haplotaxida Michaelsen, 1900**

**Family Hormogastridae Michaelsen, 1900**

**Genus Carpetania Marchán, Fernández, Díaz Cosín & Novo, 2018**

#### Description

##### External characters

Average number of segments from 203 to 323. Average weight from 1.22 g to 7.6 grams. Clitellum in segments (12)13-27. Tubercula pubertatis in segments 22-25. Pigmentation absent, color: greyish-fleshy. Cephalic keels present, moderately developed. No lateral expansions of the clitellum. Chaetae disposition geminate. No genital papillae in cd. Posterior genital papillae constrained within the extension of the clitellum. Cephalic segments not imbricated.

##### Internal characters

First thickened septum in 6/7. Last thickened septum in 9/10. Backward displacement of dorsal insertion of septum 9/10, one or two segments. No forward displacement of dorsal insertion of septa 7/8, 8/9. Spermathecal pores in intersegments 9/10, 10/11. Spermathecae tubular, the first pair smaller, with no repetition. Seven pairs of clearly developed hearts. Three oesophageal gizzards in segments 6, 7 and 8. Five typhlosole lamellae. Genital chaetae lanceolate, with strong dorsoventral differentiation and tip ornamentation; pore present, teeth present, dorsal depression absent, ventral groves absent. First nephridia with caeca between segments 10 and 12

### *Names of the species are placeholders to avoid taxonomic conflicts once the manuscript is published

*Carpetania* **species B\*** Álvarez 1977

#### Type material

Holotype and paratypes - 12 adult and subadult individuals collected in Siguero, Segovia, deposited by J. Álvarez in the Spanish Entomology Institute collection with numbers 4661-4669 and 46610-46612. Topotypes: 8 adults (UCMLT 00368-00375), 41.185 −3.6186, from a meadow in the outskirts of the village of Siguero, Segovia (Spain), collectors Darío J. Díaz Cosín, Marta Novo, Dolores Trigo.

#### Distribution

Northernmost Community of Madrid, Southern Segovia, Southern Soria (Fig. 4). Full list of known localities are shown in Suppl. table 2.

**Figure 4.**
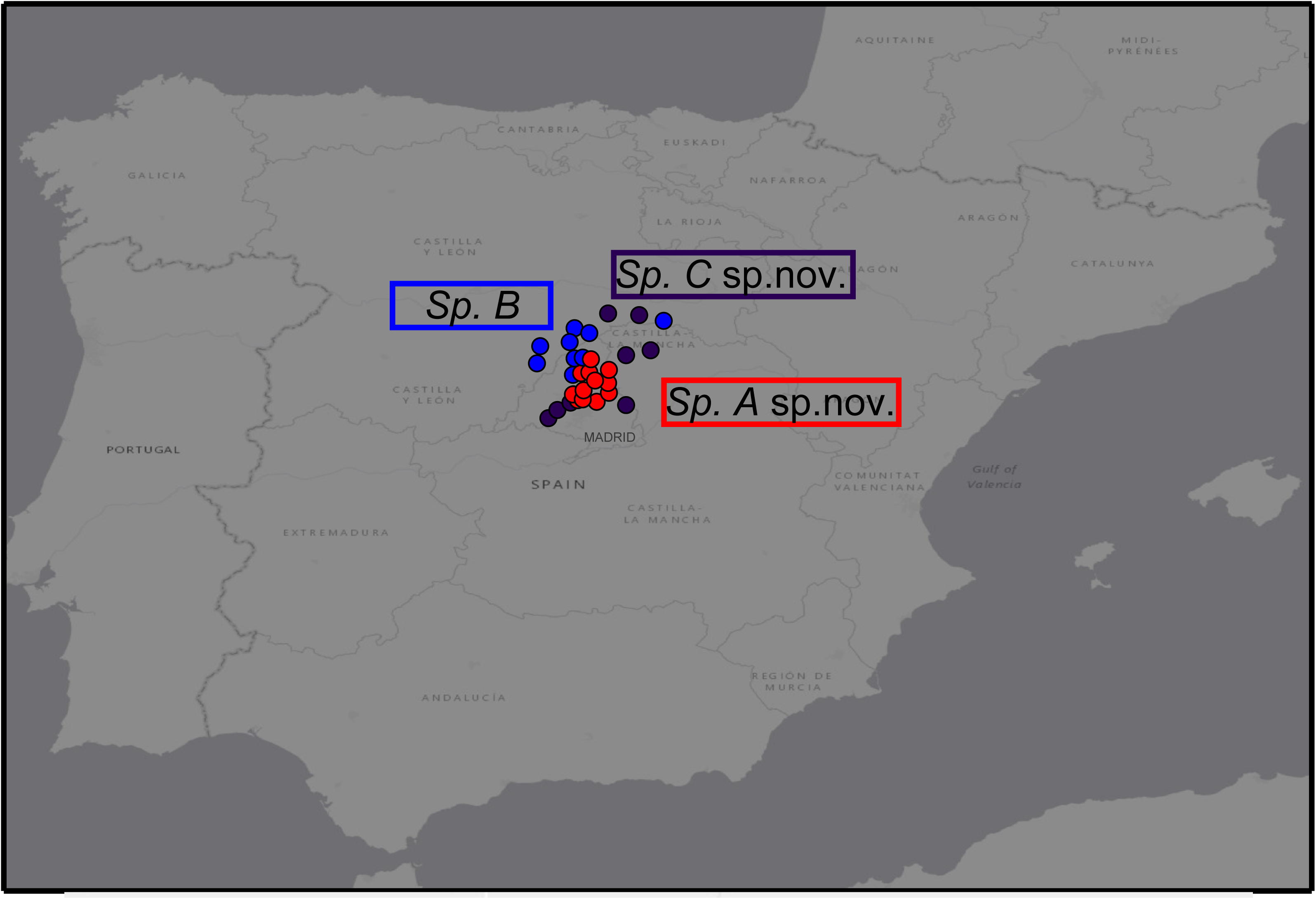
Known distribution of the three species of *Carpetania*.

#### Phylogenetic definition

includes the common ancestor of all the populations assigned in Marchán et al. (submitted) to the cluster B, and all its descendants. Known populations are shown in Suppl. table 2.

#### Reference sequences

COI - accession EF653893.1; 16S-tRNAs-accession GQ409710.1; 28S - accession GQ409654.1; H3 - accession HQ622033.1.

#### Description

External and internal morphological characters match the description of the genus *Carpetania* in all aspects but the following.

Average weight from 1.87 to 7.44 (mean= 4.98). Average number of segments from 231 to 288 (mean= 248). Generally heavier and shorter than the other species: “stout” appearance after fixation. Both pairs of spermathecae not separated by septum 9/10.

#### Diagnosis

Genital chaetae with an elongated, wide tip (Fig. 5). Diagnostic nucleotidic positions shown in table 1.

**Figure 5.**
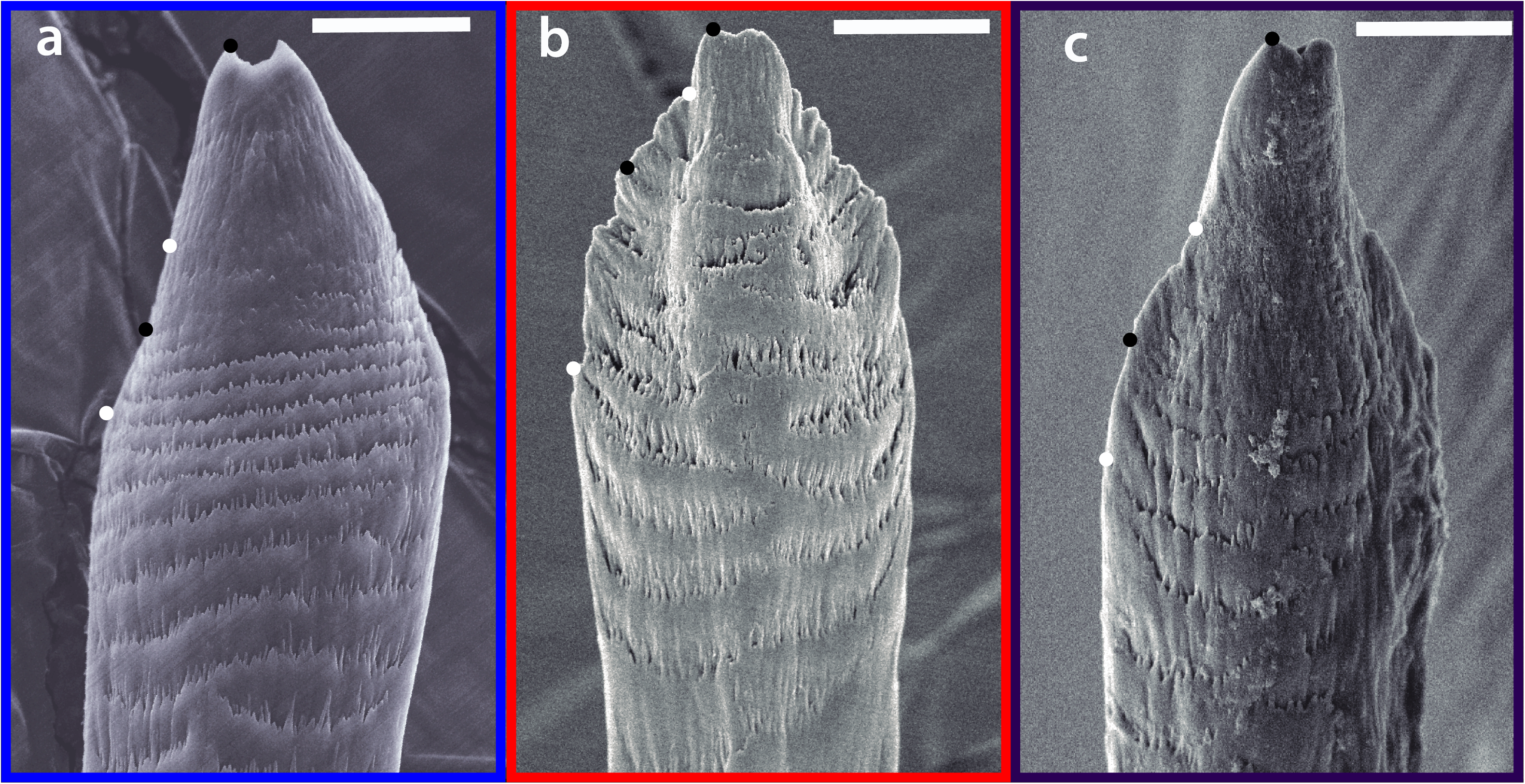
Genital chaetae distal end from representatives of the three species in *Carpetania*: a) *Carpetania* **species B**, b) *Carpetania* **species A**, c) *Carpetania* **species C**. Black and white dots show the landmarks used in geometric morphometrics analyses for reference. Scale bar: 10 μm.

#### Remarks

this species includes specimens from Siguero, topotypes of the holotype of *Hormogaster elisae*. Thus, the whole species must carry this name and be the type species of the genus *Carpetania*.

*Carpetania* **species A**\* Marchán, Novo, Férnandez and Díaz Cosín sp. nov.

#### Type material

Holotype. Adult (UCMLT 00386), 40.7394 −3.5647, from a meadow in the outskirts of the village of El Molar, Madrid (Spain), collectors Marta Novo, Darío J. Díaz Cosín. Paratypes. 7 adults (UCMLT 00387-00393), with the same collection data of the holotype

#### Distribution

Northern and Central Community of Madrid, Madrid-Guadalajara border (Fig. 4). Full list of known localities are shown in Suppl. table 2.

#### Phylogenetic definition

includes the common ancestor of all the populations assigned in Marchán et al. (submitted) to the cluster A, and all its descendants. Known populations are shown in Suppl. table 2.

#### Reference sequences

COI - accession EF653876.1; 16S-tRNAs-accession JN209295.1; 28S - accession GQ409653.1; H3 - accession JN209636.1;

#### Description

External and internal morphological characters match the description of the genus Carpetania in all aspects but the following.

Average weight from 1.22 to 4.66 (mean= 3.10). Average number of segments from 203 to 285 (mean= 253). “Slender” appearance after fixation. Both pairs of spermathecae separated by septum 9/10 in most populations (except in basal clades).

#### Diagnosis

Genital chaetae with a shortened tip and serrated lateral ridges (Fig. 5). Diagnostic nucleotidic positions shown in table 1.

*Carpetania* **species C\*** Marchán, Novo, Fernández and Díaz Cosín sp. nov.

#### Type material

Holotype. Adult (UCMLT 00376), 40.4306 −3.925, from a open holm oak woodland in the outskirts of the village of Boadilla del Monte, Madrid (Spain), collectors Marta Novo, Darío J. Díaz Cosín. Paratypes. 9 adults (UCMLT 00377-00385) with the same collection data of the holotype

#### Distribution

Southern and Central Community of Madrid, Northeastern Segovia, Southwestern Soria, Northwestern Guadalajara (Fig. 4). Full list of known localities are shown in Suppl. table 2.

#### Phylogenetic definition

includes the common ancestor of all the populations assigned in Marchán et al. (submitted) to the cluster C, and all its descendants. Known populations are shown in Suppl. table 2.

#### Reference sequences

COI - accession GQ409664.1; 16S-tRNAs-accession GQ409704.1; 28S - accession GQ409656.1; H3 - accession HQ622004.1;

#### Description

External and internal morphological characters match the description of the genus Carpetania in all aspects but the following.

Average weight from 1.41 to 5.60 (mean= 3.9). Average number of segments from 204 to 323 (mean= 273). Generally longer than the other species: “slender” appearance after fixation. Both pairs of spermathecae separated (or not) by septum 9/10, with variation between its internal lineages.

#### Diagnosis

Genital chaetae with elongated, narrow tip (Fig. 5). Diagnostic nucleotidic positions shown in table 1.

#### Remarks

this species includes three deeply divergent lineages. Further research on their range, ecology and other characters could merit their recognition as subspecies.

### -Recommendations for species identification within *Carpetania*

Morphological study of traits and character states included in the descriptions and diagnoses of the different *Carpetania* species allow a preliminary approximation to assignment of individuals. However, there is overlap between the range of morphometric characters, and scanning electron microscopy imaging of genital chaetae is demanding for untrained researchers.

The most straightforward and effective approach to species identification would consist on sequencing one or more of the proposed reference molecular markers (COI, 16S-tRNAs, 28S, H3) and their comparison with the reference sequences provided in the descriptions. This would provide a reliable assignment to one of the species.

This method does not require expertise in earthworm taxonomy and should facilitate the inclusion of these species into community and soil ecology studies. Considering existing evidence of different ecological preferences between *Carpetania* species (personal communication), it is necessary that they are not bunched together in such analyses. It is also a requisite to evaluate the conservation status of the pseudocryptic species and to preserve the diversity of this endemic genus through conservation actions.

## Acknowledgments

This work was supported by Universidad Complutense de Madrid and Santander Group under grant Proyecto de Investigación Santander/Complutense PR41/17-21027; Systematics Research Fund (SRF) and Xunta de Galicia. Consellería de Cultura, Educación e Ordenación Universitaria. Secretaria Xeral de Universidades under grant ED431B 2019/038.

RF was funded by a Marie Skłodowska-Curie Fellowship (747607). DF was funded by a Juan de La Cierva-Formación grant (FJCI-2017-32895) from the Spanish Ministry of Sciences, Innovation and Universities. MN was funded by a Postdoctoral Fellowship UCM.

